# Associations between ADHD symptom remission and white matter microstructure: a longitudinal analysis

**DOI:** 10.1101/2020.09.24.311654

**Authors:** A.E.M. Leenders, C.G. Damatac, S. Soheili-Nezhad, R.J.M. Chauvin, M.J.J. Mennes, M.P. Zwiers, D. vanRooij, S.E.A. Akkermans, J. Naaijen, B. Franke, J.K. Buitelaar, C.F. Beckmann, E. Sprooten

## Abstract

**Background:** Attention-deficit hyperactivity disorder (ADHD) is associated with white matter (WM) microstructure. Our objective was to investigate how WM microstructure is longitudinally related to symptom remission in adolescents and young adults with ADHD.

**Methods:** We obtained diffusion-weighted imaging (DWI) data from 99 participants at two time points (mean age baseline: 16.91 years, mean age follow-up: 20.57 years). We used voxel-wise Tract-Based Spatial Statistics (TBSS) with permutation-based inference to investigate associations of inattention (IA) and hyperactivity-impulsivity (HI) symptom change with fractional anisotropy (FA) at baseline, follow-up, and change between time points.

**Results:** Remission of combined HI and IA symptoms was significantly associated with reduced FA at follow-up in the left superior longitudinal fasciculus and the left corticospinal tract (CST) (*P_FWE_*=0.038 and *P_FWE_*=0.044, respectively), mainly driven by an association between HI remission and follow-up CST FA (*P_FWE_*=0.049). There was no significant association of combined symptom decrease with FA at baseline or with changes in FA between the two assessments.

**Conclusions:** In this longitudinal DWI study of ADHD using dimensional symptom scores, we show that greater symptom decrease is associated with lower follow-up FA in specific WM tracts. Altered FA thus may appear to follow, rather than precede, changes in symptom remission. Our findings indicate divergent WM developmental trajectories between individuals with persistent and remittent ADHD, and support the role of prefrontal and sensorimotor tracts in the remission of ADHD.

## Introduction

Attention-deficit/hyperactivity disorder (ADHD) is a common neuropsychiatric disorder characterized by developmentally inappropriate levels of inattention (IA) and/or hyperactivity-impulsivity (HI), with an estimated prevalence of 5% in children and adolescents and 2.5% in adults(Faraone et al., 2015). For many, ADHD begins in childhood, but the long-term clinical course of ADHD varies widely between individuals(American Psychiatric Association, 2013). Prospective studies suggest that although only 15% of children with ADHD continue to fully meet diagnostic criteria in adulthood, 60-70% of them retain impairing symptoms in adulthood(Faraone et al., 2015). ADHD diagnosis has been associated with altered patterns of brain structure and function, however the neural mechanisms related to symptom progression (i.e. remission vs. persistence) have not yet been fully unraveled (Aoki, Cortese, & Castellanos, 2018; Damatac et al., 2020; Francx, Zwiers, et al., 2015; Franke et al., 2018; Hoogman et al., 2017). Understanding this could help develop and tailor treatments to benefit long-term outcomes for children with ADHD.

The underlying neural mechanisms that drive symptom remission may be distinct from those that drive ADHD onset, thus the brains of remitted individuals could be methodologically differentiated from those of people who were never diagnosed with ADHD(Halperin & Schulz, 2006). Here, we refer to symptom remission as a dimensional concept, as a decrease in symptom severity between two time-points. Symptom remission can be driven by a number of neurodevelopmental mechanisms which are not mutually exclusive. Previous hypotheses suggest that disorder onset is characterized by a fixed anomaly or ‘scar’, while symptom remission or persistence is associated with brain maturation and normalization, or compensation and reorganization(Sudre, Mangalmurti, & Shaw, 2018). The trajectories of remission and persistence from childhood through adulthood occur in parallel to or in interaction with other neurodevelopmental processes (e.g.development of executive functions).

The development of frontal and temporal areas engaged in emotional and cognitive processes does not plateau until adulthood, which coincides with the typical age range of ADHD symptom remission(Faraone et al., 2015; Lebel & Deoni, 2018). Maturation in these regions may compensate for the initial childhood development of ADHD symptoms through top-down regulatory processes, leading to eventual symptom remission(Halperin & Schulz, 2006). Therefore, longitudinal cohort studies are essential to dissect the upstream, parallel, or downstream brain mechanisms in reference to symptom remission. Compared to a cross-sectional approach, a longitudinal design provides not only unique insights into the temporal dynamics of underlying biological processes, but also increased statistical power by reducing between-subject variability(Madhyastha et al., 2014).

Neurodevelopmental mechanisms underlying the variable long-term course of ADHD may be partly traceable using neuroimaging. Healthy brain development has been characterized using structural and functional magnetic resonance imaging (MRI), showing trajectories across the lifespan of regional volumes and activity/connectivity, respectively(Brouwer et al., 2020; van Duijvenvoorde, Westhoff, de Vos, Wierenga, & Crone, 2019). Those that have applied diffusion, magnetization transfer, relaxometry, and myelin water imaging methods have also demonstrated consistent, rapid white matter (WM) microstructural changes in the first three years of life, reflecting increased myelination or axonal packing(Lebel & Deoni, 2018). These changes continue throughout adolescence and are associated with corresponding age-related changes in gray matter(Giorgio et al., 2008). However, regarding later childhood and adolescence, the paucity of congruous findings in other WM imaging modalities besides diffusion weighted imaging (DWI) suggests that changes are primarily related to myelination and axonal packing(Lebel & Deoni, 2018). With age, WM increases in overall volume, becoming more myelinated in a region-specific fashion and reaching peak values later in life(Kochunov et al., 2012; Paus et al., 2001). The rate of development differs between WM regions, progressing in an outward, central-to-peripheral direction, wherein sensory and motor regions generally mature the earliest.

DWI studies have revealed WM microstructural abnormalities in ADHD, specifically using fractional anisotropy (FA), which is the metric we focus on here(Aoki et al., 2018; Damatac et al., 2020; van Ewijk et al., 2014; Francx, Oldehinkel, et al., 2015; Shaw et al., 2015; Sudre et al., 2018). DWI reveals information about anatomical connectivity in the brain *in vivo* by measuring the directionality of water diffusion in WM tracts, thus enabling inferences about underlying brain mechanisms by quantifying associated changes in (inter)cellular space(Beaulieu, 2002; van Ewijk et al., 2014). FA is an indirect measure of microstructural integrity—sensitive to myelination, parallel organization and fiber bundle density. A systematic meta-analysis of case-control DWI studies in ADHD found that lower FA in ADHD has mostly been reported in interhemispheric, frontal and temporal regions— however, higher FA has also been found in similar areas(Aoki et al., 2018). Given these previous WM associations with ADHD and the brain’s maturation in those same areas during an age range typical for symptom remission, the next step is to determine how WM microstructural alterations coincide with remission versus persistence of ADHD symptoms over time.

Not many longitudinal studies have examined the neurobiological underpinnings of symptom remission in WM—and none have longitudinally applied DWI. While there are no previous studies with longitudinal DWI measurements, there have been some clinical longitudinal studies with one DWI measurement. A follow-up DWI analysis three decades after diagnosis supports the theory of the disorder as an enduring neurobiological trait independent of remission; both remittent and persistent probands with an ADHD diagnosis in childhood had widespread reduced FA compared to those who did not have childhood ADHD(Cortese et al., 2013). Conversely, a network connectivity analysis of two clinical assessments and one resting-state functional MRI measurement at follow-up pointed to the presence of compensatory mechanisms that aid symptomatic remission in prefrontal regions and the executive control network: higher connectivity at follow-up was associated with HI decreases (Francx, Oldehinkel, et al., 2015). A study performed with a sample overlapping with the current study (but at an earlier sampling time with mean age 11.9-17.8 years) found, somewhat counterintuitively, that more HI symptom remission was associated with lower FA in the left corticospinal tract (lCST) and left superior longitudinal fasciculus (lSLF) at follow-up(Francx, Zwiers, et al., 2015). This previous study included clinical data from two time-points and only one DWI time-point.

Our current investigation is a continuation of our earlier DWI work in this cohort, and extends upon it in three ways. First, by capturing an older age range, we have a more complete picture of symptom remission (mean age range: 16.91-20.57 years; <u>Figure 1</u> illustrates how our study chronologically relates to that of Francx *et al.*(2015). Second, DWI measurements at two time-points allow for a more thorough investigation of the chronology and mechanisms of FA development in relation to symptom remission. Third, we used Permutation Analysis of Linear Models(PALM), a newly available permutation-based analysis technique, to account for the family structure in our sample(Winkler, Webster, Vidaurre, Nichols, & Smith, 2015). We aimed to examine whether symptom remission may be underpinned by WM alterations as adolescents with ADHD develop into adulthood. Given our longitudinal DWI data, we were able to distinguish between (1) pre-existing WM features that predict the likelihood of symptoms to remit or persist, (2) WM changes over time that occur concurrently with symptom change, and (3) WM alterations that may be a (direct or indirect) downstream consequence of symptom remission versus persistence.

**Figure 1.**
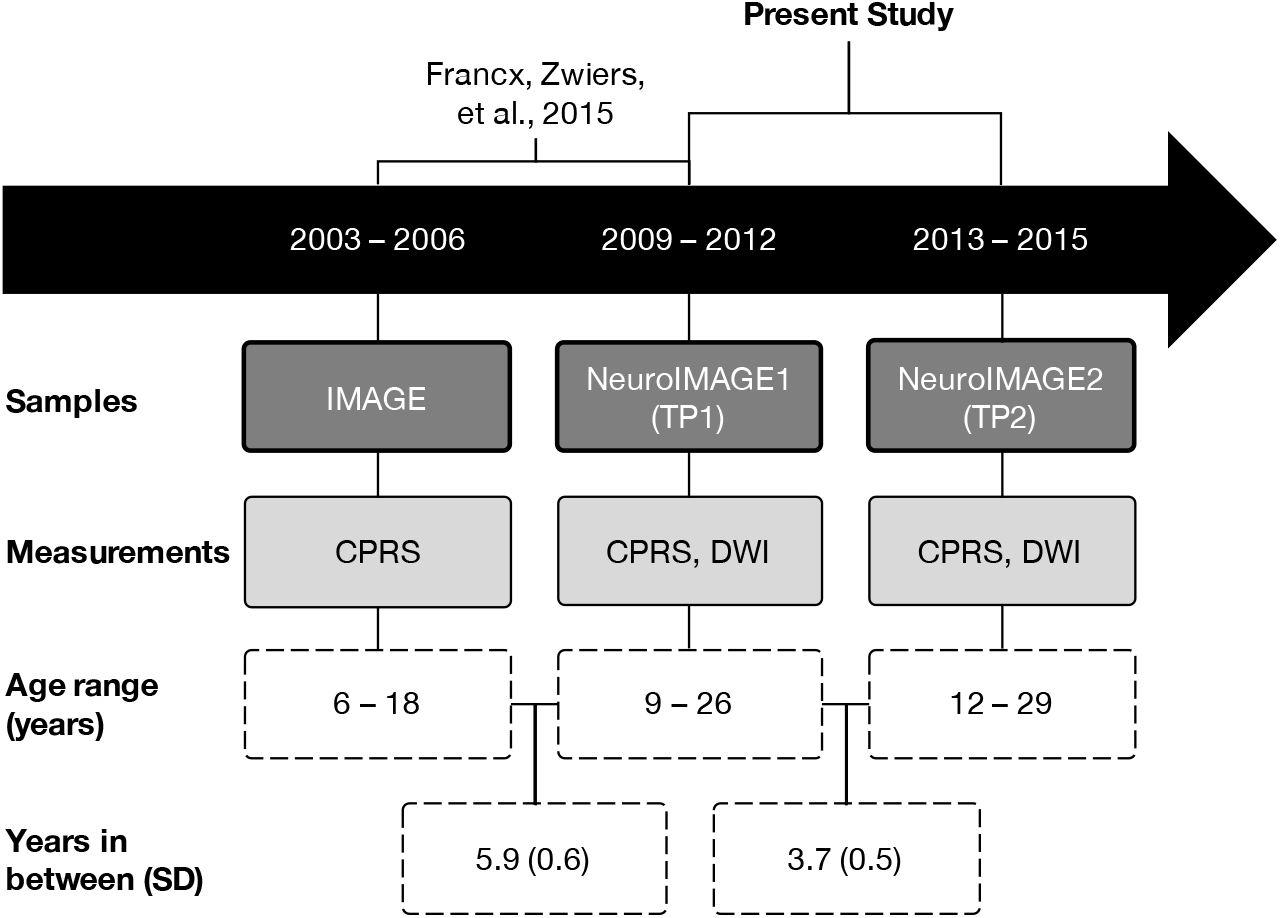
Schematic diagram of how this study chronologically relates to a previous study, samples included, relevant clinical and imaging measurements, study sample age ranges, and mean years (standard deviation) in between each measurement time-point(Francx, Zwiers, et al., 2015; Müller et al., 2011b; von Rhein et al., 2015). The present study is an analysis of TP1 and TP2. DWI: diffusion-weighted imaging, CPRS: Conners Parent Rating Scale(Conners et al., 1999).

## Methods

### Participants

Clinical and MRI data were collected in two waves from probands with childhood ADHD, their first-degree relatives, and healthy families: NeuroIMAGE1(TP1) and NeuroIMAGE2(TP2)(Müller et al., 2011a, 2011b; von Rhein et al., 2015). The current study included probands, affected and unaffected siblings, and healthy controls who participated in both TP1 and TP2 and had DWI data from both waves(N=120). After we excluded subjects(<u>Figure S1</u>), there were 99 participants from 65 families in our final sample (see characteristics in <u>Table 1</u>). For both time-point groups, there were no differences between the participants included in the current analysis and the complete sample on measures of ADHD severity, age, and sex(P>0.05). We normalized head motion z-scores after excluding outliers. Global FA at TP1, TP2, and the difference between TP1 and TP2 were normally distributed.

**Table 1.**
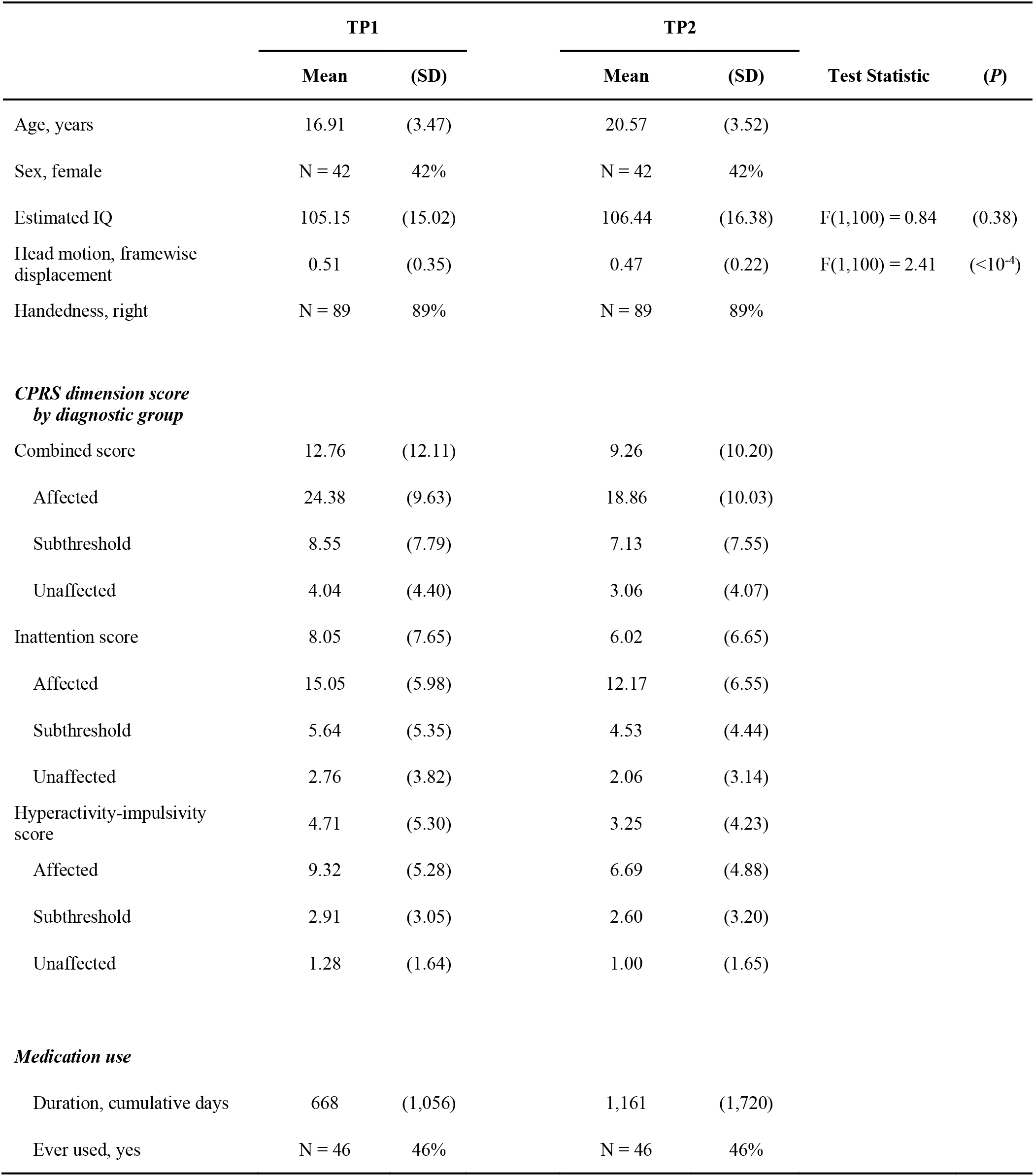
Demographic and clinical characteristics of the sample at NeuroIMAGE1(TP1) and NeuroIMAGE2(TP2) with mean and standard deviation. Reported values pertain to all participants in the final sample after all quality control(N=99). IQ: estimated at both timepoints using vocabulary and block design subtests of the Wechsler Intelligence Scale for Children(WISC-III) or Wechsler Adult Intelligence Scale(WAIS-III). Combined CPRS symptom score is the sum of two separate dimensions: hyperactivity-impulsivity and inattention. Medications: Ritalin(methylphenidate), Concerta(methylphenidate), Strattera(atomoxetine), and any other ADHD medication. The majority of patients were taking prescription medication for ADHD, mostly methylphenidate or atomoxetine. Medication use duration: lifetime cumulative number of days used on the day of the MRI scan (numerical integer value). Ever used: whether or not participants had ever taken ADHD medication in their lives (binary factor: yes or no).

Given the longitudinal design of our study, we did not split our participants into cases versus controls; there were those who were diagnosed as unaffected at both time-points, as well as those who had a symptom score of zero in all dimensions at both time-points (<u>Figure S2</u>). Our sample includes controls, or people who do not have (subthreshold) ADHD; however, this group of people changed between the waves of the study (<u>Figure S3</u>). Some individuals originally recruited as controls or unaffected siblings developed ADHD at a later time point and others recruited as patients remitted. Moreover, ADHD may be operationalized as a continuous trait rather than a binary diagnostic variable, especially in longitudinal studies (Lahey & Willcutt, 2010; Marcus & Barry, 2011). In a previous cross-sectional study we specifically showed that, compared to categorical diagnoses, continuous symptom measures were more sensitive to diffusion-weighted brain features in this sample (Damatac et al., 2020). Therefore, we focus on participants’ ADHD symptom scores in all models, thus optimizing our design and methods for capturing the dynamic and continuous nature of the ADHD spectrum throughout adolescence.

### Clinical measurements

Conners Parent Rating Scale(CPRS) questionnaires were used to assess the severity of 27 inattention(IA) and hyperactive-impulsive(HI) symptoms at TP1 and TP2(Conners et al., 1999). We used CPRS instead of the self-rated report because it was the consistent measure across waves and ages. We used raw CPRS scores to increase the distribution width, and analyzed HI, IA, and combined symptom scores per subject, per time-point. Here, we define symptom change as the score difference:ΔCPRS=CPRS_TP1_–CPRS_TP2_. Thus, a more positive Δ value indicates more symptom improvement.

### Data acquisition and DWI pre-processing

MRI data were acquired with a 1.5-Tesla AVANTO scanner(Siemens, Erlangen, Germany). The scanner was equipped with an 8-channel receive-only phased-array head coil. Whole-brain diffusion-weighted images were collected(twice refocused pulsed-gradient spin-echo EPI; 60 diffusion-weighted directions spanning an entire sphere; b-value 1000s/mm^2^; 5 non-diffusion weighted images; interleaved slice acquisition; TE/TR=97/8500ms; GRAPPA-acceleration factor 2; no partial Fourier; voxel size 2×2×2.2mm). DWI acquisition parameters are described in detail elsewhere(von Rhein et al., 2015).

### Longitudinal TBSS

We performed whole brain voxel-wise analyses with Tract-Based Spatial Statistics (TBSS)(Smith et al., 2006). Our study’s longitudinal design necessitated an analysis pipeline that considered how within-subject changes may be greater than between-subject changes; intra-subject data alignment across time brings extra difficulty compared to cross-subject nonlinear registration to common space. We used a bespoke pipeline adapted from others to create non-biased individual subject templates(Madhyastha et al., 2014). <u>Figure 2</u> summarizes our pipeline(detailed in <u>Figure S1</u>).

**Figure 2.**
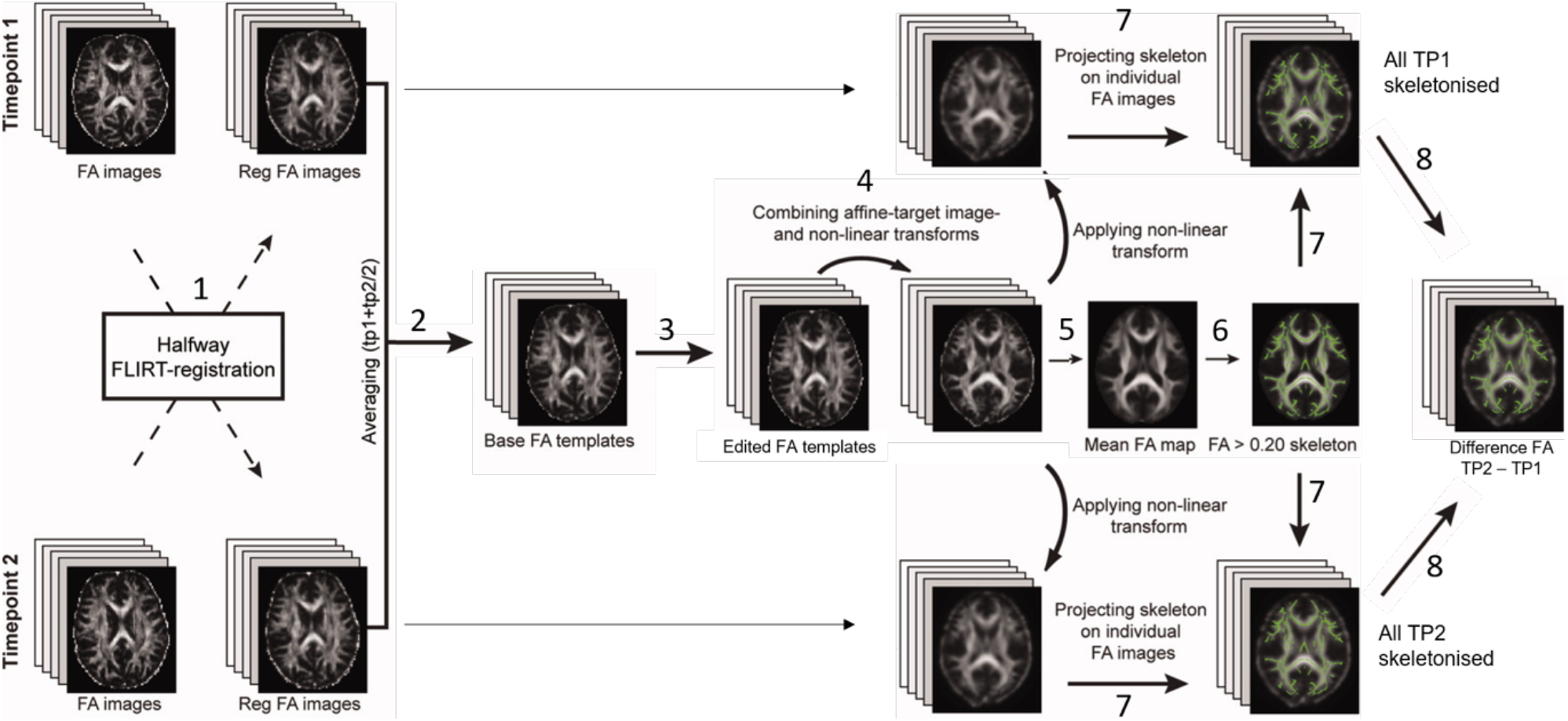
Our longitudinal TBSS pipeline was adapted to create a nonbiased individual subject template for use as a base template (2), which was then non-linearly registered to FMRIB58 FA standard-space (4), to create a mean FA skeleton (5), onto which each subject’s aligned FA data from both time points was projected (7).

### Statistical analysis

We constructed three general linear models for our voxel-wise analyses (<u>Table S1</u>). We kept difference in raw CPRS score(ΔCPRS) as a constant predictor, while separately testing FA at baseline(FA_TP1_), follow-up(FA_TP2_), and the difference between TP1 and TP2 (ΔFA) as dependent variables. Our covariates included sex, normalized head motion(framewise displacement) at respective time point(s), age at TP1, age difference between TP1 and TP2, and CPRS symptom score at TP1(<u>Table S1</u>, <u>Figure S4</u>). Our main analyses first examined combined symptom scores and, if significant, subsequent analyses examined whether effects were driven by HI or IA.

We used PALM to account for the lack of independence in the data due to sibling relationships and shared variance between families, constraining permutation tests between families of the same sizes(Winkler, Ridgway, Webster, Smith, & Nichols, 2014). We designed multi-level exchangeability blocks which did not allow permutation among all individuals; permutations were constrained both at the whole-block level (i.e. permute between families of the same size) and the within-block level (i.e. permute within families)(<u>Figure S5</u>). We corrected for multiple testing by running 5000 permutations and threshold-free cluster enhancement(TFCE) as implemented in PALM, part of the FSL toolbox (https://fsl.fmrib.ox.ac.uk/fsl/fslwiki/PALM)(Smith & Nichols, 2009). Results with TFCE-corrected *P*<0.05 were considered statistically significant. All tests used the standard parameter settings for height, extent, and connectivity: H=2, E=1, C=26. We used the Johns Hopkins University DTI-based WM atlas (https://fsl.fmrib.ox.ac.uk/fsl/fslwiki/Atlases) to relate significant clusters to known WM tracts.

## Results

### Symptom change over time

In <u>Table 1</u>, we present mean symptom scores for HI, IA, and combined (HI+IA). Combined symptom scores significantly decreased over time (*t*(98)=4.884,*P_FWE_*=2.027×10^−6^). This was due to decreases in both IA scores (*t*(98)=4.226,*P_FWE_*=2.672×10^−5^) and HI scores (*t*(98)=4.394,*P_FWE_*=1.410×10^−5^), with a mean decrease of 2.04(SD=4.80) in IA, and 1.46(SD=3.30) in HI score.

### Symptom change in relation to WM microstructure at two time-points

As a point of reference: Participants who showed relatively more symptom remission had FA values that moved neither towards nor away from that of those who were diagnosed as never affected; their global FA values increased over time at a similar rate as the never affected individuals, while those who essentially persisted or developed more symptoms over time showed a slightly steeper increase in FA than the other two groups (<u>Figure S6</u>).

Our models demonstrated no significant association between combined symptom score remission and FA_TP1_, but there was a significant negative association between combined symptom score remission and FA_TP2_ in two regions according to the atlas: lSLF(*P_FWE_*=0.038) and lCST(*P_FWE_*=0.044)(<u>Table 2</u>;<u>Figure 3A</u>). This was mainly driven by a negative effect of HI dimension score on FA in lCST(*P_FWE_*=0.049)(<u>Table 2</u>;<u>Figure 3B</u>).

**Table 2.** TBSS results of significant models: WM tracts, peak voxels, and localization of significant clusters(*P_FWE_*<0.05) of voxel-wise permutation based dimensional analyses (see full composition of models in Table S1). N_voxels_: number of voxels, X/Y/Z_COG_: location of the center of gravity for the cluster(vox/mm), MNI: Montreal Neurological Institute coordinates, t_max_: highest threshold-free cluster enhancement t-statistic value per cluster, WM tract: anatomical location in a white matter tract based on the Johns Hopkins University DTI-based white-matter atlases, lCST: left corticospinal tract, lSLF: left superior longitudinal fasciculus. Model FA_TP2_ ~ ∆CPRS_combined_: Less combined symptom score decrease was associated with more FA at follow-up in lSLF and lCST. Model FA_TP2_ ~ ∆CPRS_HI_: The negative effect in the previous model was driven by HI score remission.

| Model | WM tract | N<sub>voxels</sub> | MNI (Peak voxel) |  |  | *t<sub>max</sub>* | *P<sub>FWE</sub>* |
| --- | --- | --- | --- | --- | --- | --- | --- |
|  |  |  | X<sub>COG</sub> | Y<sub>COG</sub> | Z<sub>COG</sub> |  |  |
| FA<sub>TP2</sub> ~ ΔCPRS<sub>combined</sub> | ICST | 723 | -21 | -27 | 44 | 0.755 | **0.044** |
|  | ISLF | 579 | -33 | -20 | 38 | 0.881 | **0.038** |
| FA<sub>TP2</sub> ~ ΔCPRS<sub>HI</sub> | ICST | 17 | -18 | -25 | 52 | 0.981 | **0.049** |

**Figure 3.**
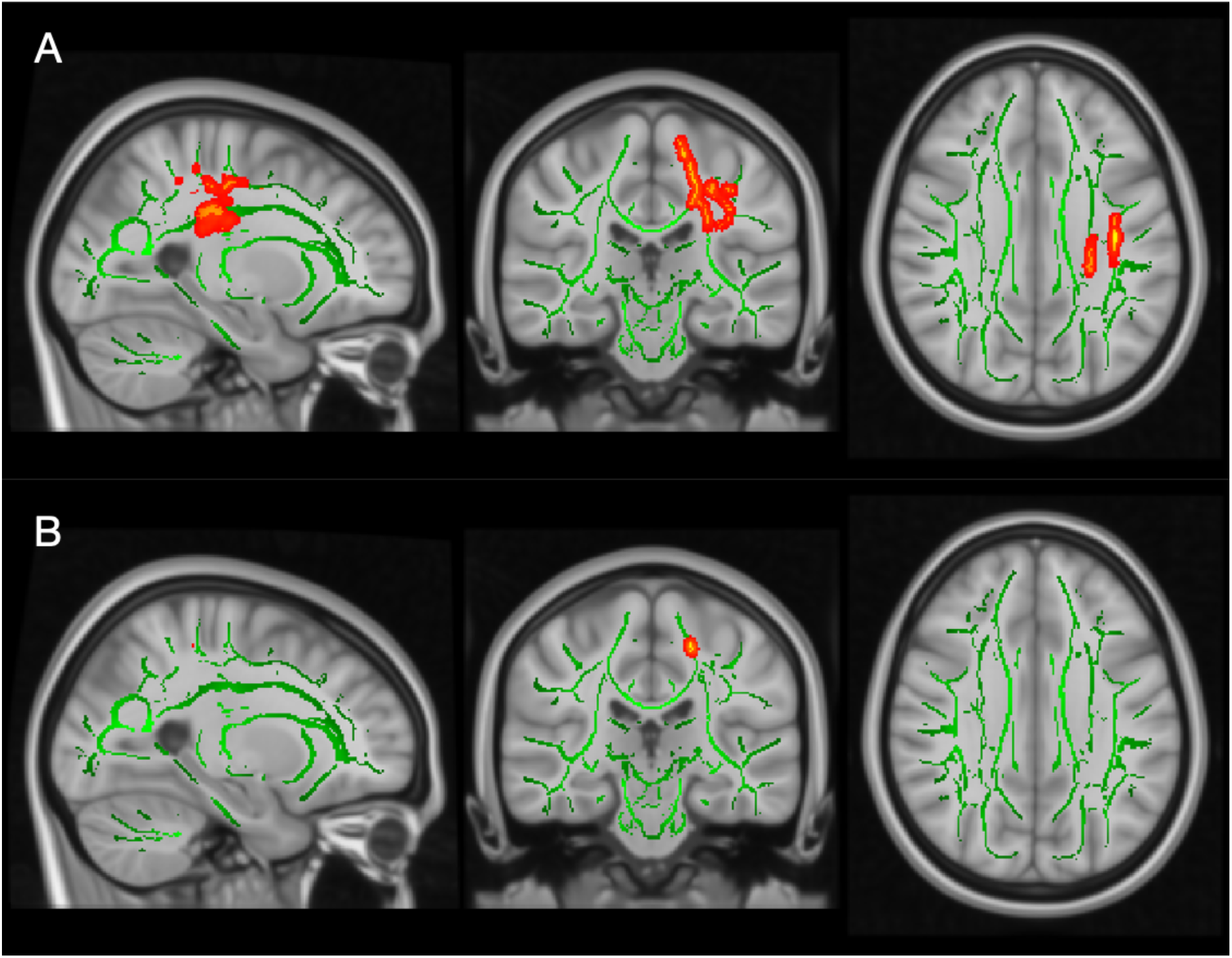
TBSS results showing significant associations (red-yellow) between FA values and the CPRS dimension scores over time. The mean FA skeleton across all subjects (green) was overlain on the MNI template image for presentation (x=−25, y=−25, z=31). Results were thickened for visualization (FSL “tbss_fill”) and presented here in radiological convention from sagittal, coronal, and axial perspectives, respectively. **(A)**Lower FA values at follow-up (TP2) were associated with a larger decrease in combined symptom score in the left superior longitudinal fasciculus (lSLF)(*P_FWE_*=0.038) and the left corticospinal tract (lCST)(*P_FWE_*=0.044); **(B)**Lower FA values at TP2 were associated with a larger decrease in HI symptom score in lCST(*P_FWE_*=0.049).

Additionally, there was a negative trend association between combined symptom score difference and FA difference (*P_FWE_*=0.079). Because our one model with a significant effect, as well as those previously reported in an overlapping sample at an earlier time window, were driven by HI, we performed an exploratory *post-hoc* analysis on symptom score difference and FA difference with only HI dimension scores(Brouwer et al., 2020; Damatac et al., 2020; Francx, Zwiers, et al., 2015). Our exploratory results show that larger HI symptom decrease was associated with a larger decrease in FA over time in ten clusters spread over six WM tracts(<u>Table S2</u>, <u>Figure S7</u>).

### Post hoc tests of confounders and demographics

Sex, normalized head motion at respective time point(s), age at TP1, age difference between TP1 and TP2, and CPRS symptom score at TP1 were included as covariates in all models. We report main effects after the removal of non-significant interaction effects in <u>Table 2</u>. We found neither a significant main effect nor an interaction effect of any of these covariates with ΔCPRS for all analyses reported above(<u>Table S3</u>).

## Discussion

In this longitudinal investigation of ADHD and WM microstructure, we report that higher symptom decrease is associated with lower FA at follow-up in lCST and lSLF, an effect mainly driven by HI symptom decrease. Thereby, we have essentially replicated and extended the findings reported by Francx, Zwiers, et al. (2015) at an older age range, contributing to the growing body of evidence describing the progression of ADHD and its relation with WM. Additionally, we utilized an improved statistical method to account for the family structure in our data, thus confirming that previous results in this cohort were not confounded by within-family correlations(Winkler et al., 2015). By substantiating those earlier findings, with replication in participants at an older age, and upon better accounting for family relatedness, we conclude that a decrease in symptoms from early adolescence is associated with lower FA in late adolescence and young adulthood.

Our longitudinal design of two diagnostic and two DWI time-points allows us to speculate about the chronology of brain changes versus symptom changes. First, we found no evidence that baseline FA predicts ADHD symptom change over time. Second, though a natural expectation would be that more remission leads to higher FA, we found the opposite, somewhat paradoxical result: Greater ADHD symptom decrease was associated with lower FA at follow-up in lSLF and lCST. Third, we found that HI, not IA, symptom decrease was the main driver behind the association with reduced FA in lCST. WM microstructure can change in response to behavior or learning (i.e. plasticity). It is possible that decreased (motor) hyperactivity is associated with less use of corticospinal and motor tracts, which may lead to decreased FA in this area at TP2. Overall, lower FA in both tracts appears to follow, rather than precede, symptom decrease. Speculatively, this suggests that the WM changes may be a downstream result, rather than a cause, of symptom remission in ADHD.

FA is an indirect reflection of microstructure and some neuronal processes that improve anatomical connectivity may paradoxically manifest as decreased FA in some locations— especially in principal WM highways through which several fibers cross, like the SLF and CST. At the axonal level, more sprouting, pruning, crossing fibers or fiber dispersion in those tracts during maturation may demonstrate as reduced FA over time. Plasticity in myelin or axon integrity in less dominant fibers could also exhibit as reduced FA in voxels containing multiple fiber orientations. In our participants whose symptoms persisted, higher FA could be the outcome of brain reorganization in less dominant fiber tracts, particularly in those that traverse the CST and SLF. Event-related and resting-state functional MRI studies that grouped their subjects categorically have reported that remitters have stronger connectivity than persisters(Clerkin et al., 2013; Francx, Oldehinkel, et al., 2015). Lower functional connectivity in certain tracts may be related to higher FA in other tracts and vice versa. In a top-down fashion, remitters may learn compensatory strategies to overcome their ADHD symptoms as they age, while persisters may either learn disadvantageous strategies, other beneficial compensatory strategies, or none at all, leading to divergent trajectories of WM development in various brain regions in individuals with persistent ADHD symptoms(Wetterling et al., 2015).

One can find in the literature several instances wherein the SLF and CST are implicated in ADHD. The SLF generally subserves a wide variety of functions related to language, attention, memory, emotion, and visuospatial function; many studies have pointed to its function in visuospatial awareness, as well as attentional selection of sensory content(Conner et al., 2018; Shaw et al., 2015; Wolfers et al., 2015). Our findings are partly in accordance with those of others that have found neurodevelopmental effects linked with unilaterally compromised lSLF maturation during adolescence(Peters et al., 2012). Thus, ADHD symptom persistence may influence higher unilateral SLF integrity as a person develops from early adolescence to young adulthood. The CST integrates cortical and lower brain processing centers in the motor system, has an important role in modulating sensory information, and may be particularly relevant to motor hyperactivity in ADHD(Moreno-López, Olivares-Moreno, Cordero-Erausquin, & Rojas-Piloni, 2016). Altered modulation of sensory information could potentially be involved in HI remission, as the CST contains fibers running from the primary motor, premotor, supplementary motor, somatosensory, parietal and cingulate cortex to the spine and is thus involved in the control of complex voluntary distal movements(Welniarz, Dusart, & Roze, 2017). Correspondingly, the persistence of HI could, indeed, result in increased FA in CST through time. Our unilateral findings may have risen from the fact that 88% of our subjects were right-handed, and most CST axons cross to the contralateral side at the pyramidal decussation before reaching lower motor neurons(Welniarz et al., 2017). We found no evidence that handedness was correlated with change in symptom scores (<u>Table S4</u>).

Based on previous investigations that have similarly found effects of ADHD symptoms on WM microstructure driven by HI, we also conducted an exploratory analysis of only HI symptom remission effects on change in FA(Damatac et al., 2020; Francx, Zwiers, et al., 2015). Our results may suggest that HI symptom remission is associated with more decrease in FA over time. Most of the associations we found were clustered in prefrontal and frontostriatal regions. Higher functional connectivity in prefrontal networks in young adults has been associated with more improvement in symptoms over time(Clerkin et al., 2013; Francx, Zwiers, et al., 2015). Likewise, the prefrontal cortex and its connections are especially important in the remission or persistence of ADHD symptoms(Francx, Oldehinkel, et al., 2015; Halperin & Schulz, 2006). As it continues to develop throughout adolescence, the prefrontal cortex can potentially compensate for the initial causes of ADHD through its connectivity with subcortical regions such as the striatum. Indeed, a study using independent component analysis demonstrated that ADHD diagnosis was significantly associated with reduced brain volume in a component that mapped to the frontal lobes, the striatum, and their interconnecting WM tracts(Cupertino et al., 2019). Although exploratory and tentative, our finding of decreased FA in frontostriatal regions coinciding with HI symptom remission is thus in line with Halperin & Schulz’s theory (2006): Neural plasticity and the development of the prefrontal cortex and interconnected neural circuits facilitate recovery over the course of development.

We used a dimensional approach in defining the ADHD phenotype, in line with our recent findings in a large overlapping cohort wherein no evidence was found for altered FA in association with categorical ADHD diagnosis(Damatac et al., 2020). Unaffected participants clustered at the low end of the score distribution. Given the relatively small number of fully remitted patients (N=5), together with a subset of ‘partly remitted’ individuals, our use of symptom severity as a continuous variable maximized power to detect symptom-related changes, while also circumventing arbitrary decisions on the definition of remission(Du Rietz et al., 2016). We thus interpret our findings in terms of symptom severity, reflecting the degree of remission in ADHD patients as well as variation in individuals who do not reach diagnostic threshold.

Head motion is quite typical in the ADHD population and is hence a common confound in such studies(Aoki et al., 2018; Yendiki, Koldewyn, Kakunoori, Kanwisher, & Fischl, 2014; Zwiers, 2010). A previous meta-analysis of DWI studies in ADHD found that most investigations that controlled for head motion did not have significant results(Aoki et al., 2018). We normalized head motion and included it as a confound covariate in all of our analyses, as well as checked each model for an interaction effect with the head motion parameter. We found no evidence that it had an influence on our results.

FA estimates can be less accurate in brain regions consisting of so-called “kissing” and/or crossing fibers, like the CST and SLF. FA gives only one value for the overall restriction of anisotropy in a voxel, which could be a crucial aspect in the inconsistency of findings in the literature regarding WM and ADHD. Future studies may include complementary longitudinal region-of-interest tractography analyses in the clusters that we found to be significant, or by using DWI methods that deliver greater resolution at the neurite level. Techniques that utilize orientation dispersion indices or WM fiber density could potentially provide clarity in the constant discourse of how crossing fibers can mar inferences about FA and brain effects of ADHD. Likewise, incorporating additional DWI data from more than two time points throughout development would, naturally, increase statistical power and enhance our understanding of the dynamic interplay between disorder and development. Due to the wide age range in our sample, we checked for but did not find interactions with baseline age. Nonetheless, more complex, nonlinear patterns with age may exist across the age ranges we studied. We cannot assume that the relationship between symptom change and brain change is a constant process throughout the entire age range in this study.

## Conclusion

We used two DWI time-points in a longitudinal study of dimensional symptom scores in ADHD. Our results indicate that, in specific WM tracts, greater symptom improvement results in lower FA at follow-up. We show that WM alterations may occur downstream of symptom change. The effects we have found confirm and extend earlier findings in an overlapping sample; they indicate divergent trajectories of WM development in individuals with persistent ADHD symptoms compared to those showing remittance, and support the role of prefrontal and sensorimotor development in the symptom remission of ADHD.

## Abbreviations

(WM): white matter
(TP1): NeuroIMAGE1
(TP2): NeuroIMAGE2

## Acknowledgments

The NeuroIMAGE study was supported by National Institutes of Health Grant No. R01MH62873 (to Stephen V. Faraone), NWO Large Investment Grant No. 1750102007010 (to JKB), NWO Brain & Cognition Grant Nos. 056-13-015 (to JKB) and 433-09-242 (to JKB), ZonMW Grant No. 60-60600-97-193 (to JKB), and a matching grant from Radboud University Medical Center.

### Key points

- Attention-deficit hyperactivity disorder is associated with white matter microstructure, but little is known about how they are longitudinally related as a child develops into adulthood.
- We used voxel-wise Tract-Based Spatial Statistics with permutation-based inference to investigate how symptom score change relates to fractional anisotropy in individuals with ADHD and typically developing controls over a period of about four years.
- We provided for the first time, evidence that altered FA appears to follow, rather than precede, changes in symptom remission.
- Our findings indicate divergent white matter developmental trajectories between individuals with persistent and remittent ADHD, and support the role of prefrontal and sensorimotor tracts in the remission of ADHD.

---

**Table S1.** Composition of our three general linear models. We essentially have a cross-lagged design with fractional anisotropy (FA) as the dependent variable. The difference in CPRS (ΔCPRS=CPRS_TP1_–CPRS_TP2_) is the predictor variable in all models. For each model, CPRS score could be the inattention, hyperactivity-impulsivity, or combined score. The outcome variables of these models are, respectively: FA at baseline (TP1), FA at follow-up (TP2), and change in FA. Permutation analysis (PALM) necessitated that we kept the (TBSS output) FA image as the dependent variable. We covaried for participant baseline age, change in years of age, sex, baseline CPRS symptom score, and framewise displacement head motion at the relevant FA time-point in the dependent variable.

|  |  |
| --- | --- |
| **Model 1** | $FA_{TP1} \sim \Delta CPRS + age_{TP1} + \Delta age + sex + CPRS_{TP1} + head\ motion_{TP1}$ |
| **Model 2** | $FA_{TP2} \sim \Delta CPRS + age_{TP1} + \Delta age + sex + CPRS_{TP1} + head\ motion_{TP2}$ |
| **Model 3** | $\Delta FA \sim \Delta CPRS + age_{TP1} + \Delta age + sex + CPRS_{TP1} + head\ motion_{TP1} + head\ motion_{TP2}$ |

**Table S2.** Exploratory *post-hoc* analysis results. Greater HI symptom decrease was associated with a larger decrease in FA over time in several WM clusters. P-values reported here were not adjusted for multiple testing. lIFOF: left inferior fronto-occipital fasciculus, Fmin: forceps minor of the corpus callosum, lUNC: left uncinate fasciculus, CCG: cingulum cingulate gyrus.

| Exploratory<br>model | WM<br>tract | N<sub>voxels</sub> | MNI (Peak voxel) |  |  | *t<sub>max</sub>* | *P<sub>FWE</sub>* |
| --- | --- | --- | --- | --- | --- | --- | --- |
|  |  |  | X<sub>COG</sub> | Y<sub>COG</sub> | Z<sub>COG</sub> |  |  |
| $\Delta FA \sim \Delta CPRS_{III}$ | IIFOF | 508 | -29 | 24 | 17 | 0.441 | 0.041 |
|  | IIFOF | 376 | -17 | 31 | -10 | 0.613 | 0.035 |
|  | Fmin | 339 | -18 | 50 | 0 | 0.585 | 0.042 |
|  | IUNC | 174 | -25 | 17 | -8 | 0.564 | 0.045 |
|  | ICST | 158 | -22 | -13 | 8 | 0.562 | 0.047 |
|  | Fmin | 22 | -20 | 38 | 21 | 0.563 | 0.049 |
|  | CCG | 17 | -17 | 32 | 23 | 0.562 | 0.049 |
|  | IIFOF | 11 | -28 | 15 | -1 | 0.563 | 0.049 |
|  | Fmin | 11 | -18 | 46 | 17 | 0.566 | 0.049 |
|  | Fmin | 9 | -12 | 41 | -17 | 0.563 | 0.049 |

**Table S3.** Interaction effects of ΔCPRS and age at TP1 for significant models.

| Model | WM tract | MNI (Peak voxel) |  |  |  |  |  | age<sub>TP1</sub> interaction *P* |
| --- | --- | --- | --- | --- | --- | --- | --- | --- |
|  |  | N<sub>voxels</sub> | X<sub>COG</sub> | Y<sub>COG</sub> | Z<sub>COG</sub> | *t*<sub>max</sub> | *P*<sub>FWE</sub> |  |
| **FA<sub>TP2</sub> ~ ΔCPRS<sub>combined</sub>** | ICST | 723 | -21 | -27 | 44 | 0.755 | 0.044 | 0.463 |
|  | ISLF | 579 | -33 | -20 | 38 | 0.881 | 0.038 | 0.808 |
| **FA<sub>TP2</sub> ~ ΔCPRS<sub>III</sub>** | ICST | 17 | -18 | -25 | 52 | 0.981 | 0.049 | 0.154 |
| **ΔFA ~ ΔCPRS<sub>III</sub>** | IIFOF | 508 | -29 | 24 | 17 | 0.441 | 0.041 | 0.667 |
|  | IIFOF | 376 | -17 | 31 | -10 | 0.613 | 0.035 | 0.055 |
|  | Fmin | 339 | -18 | 50 | 0 | 0.585 | 0.042 | 0.874 |
|  | IUNC | 174 | -25 | 17 | -8 | 0.564 | 0.045 | 0.131 |
|  | ICST | 158 | -22 | -13 | 8 | 0.562 | 0.047 | 0.130 |
|  | Fmin | 22 | -20 | 38 | 21 | 0.563 | 0.049 | 0.239 |
|  | CCG | 17 | -17 | 32 | 23 | 0.562 | 0.049 | 0.238 |
|  | IIFOF | 11 | -28 | 15 | -1 | 0.563 | 0.049 | 0.238 |
|  | Fmin | 11 | -18 | 46 | 17 | 0.566 | 0.049 | 0.239 |
|  | Fmin | 9 | -12 | 41 | -17 | 0.563 | 0.049 | 0.237 |

**Table S4.** No significant correlation (Spearman’s rho) between difference in CPRS score (ΔCPRS=CPRS_TP1_– CPRS_TP2_) and right-handedness in our sample.

| ΔCPRS score | Spearman's $\rho$ | *P* |
| --- | --- | --- |
| Combined | 0.038 | 0.709 |
| Hyperactivity-impulsivity | 0.098 | 0.331 |
| Inattention | -0.007 | 0.948 |

**Figure S1.**
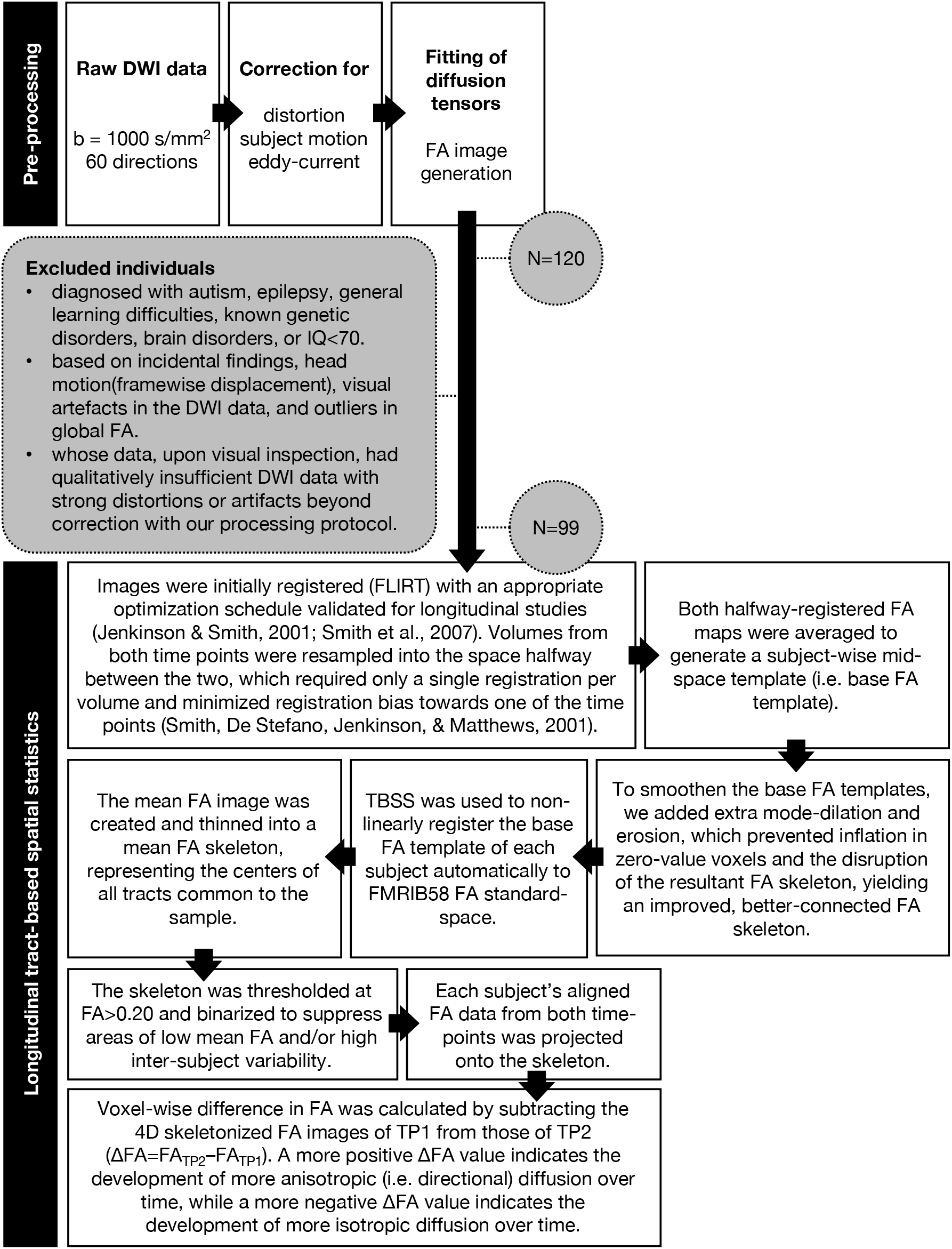
Processing pipeline. During pre-processing, DWI images were realigned and corrected for residual eddy current and for motion artefacts using robust tensor modelling (PATCH)(Zwiers, 2010). Diffusion tensor characteristics and FA values were calculated for each voxel(Behrens et al., 2003). After pre-processing, we used a custom longitudinal TBSS pipeline adapted from others to create non-biased individual subject templates

**Figure S2.**
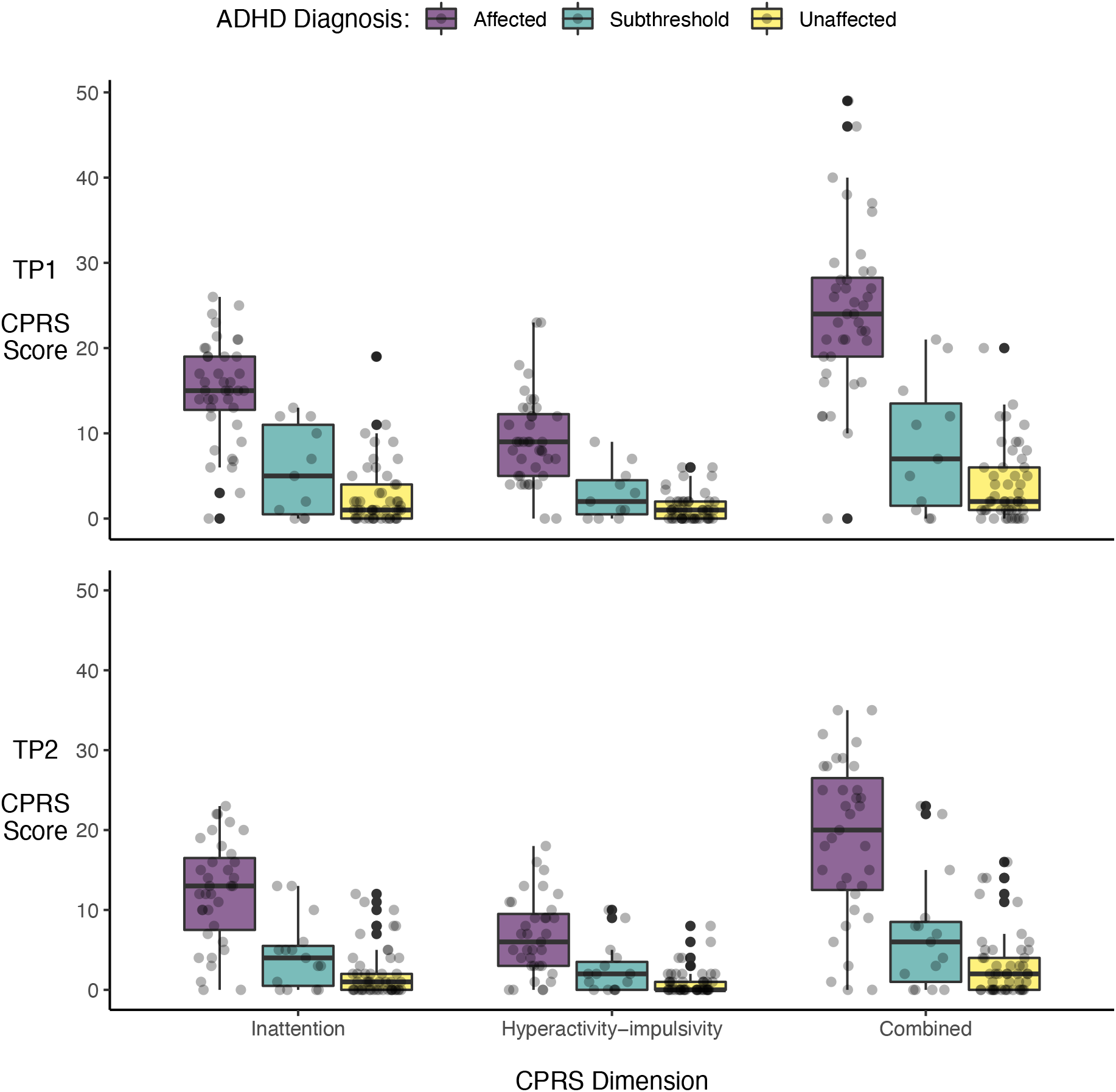
Jittered scatterplots overlain with boxplots of individual Conners Parent Rating Scale (CPRS) dimension scores colored by clinical diagnosis group at baseline (TP1; top panel) and follow-up (TP2; bottom panel).

**Figure S3.**
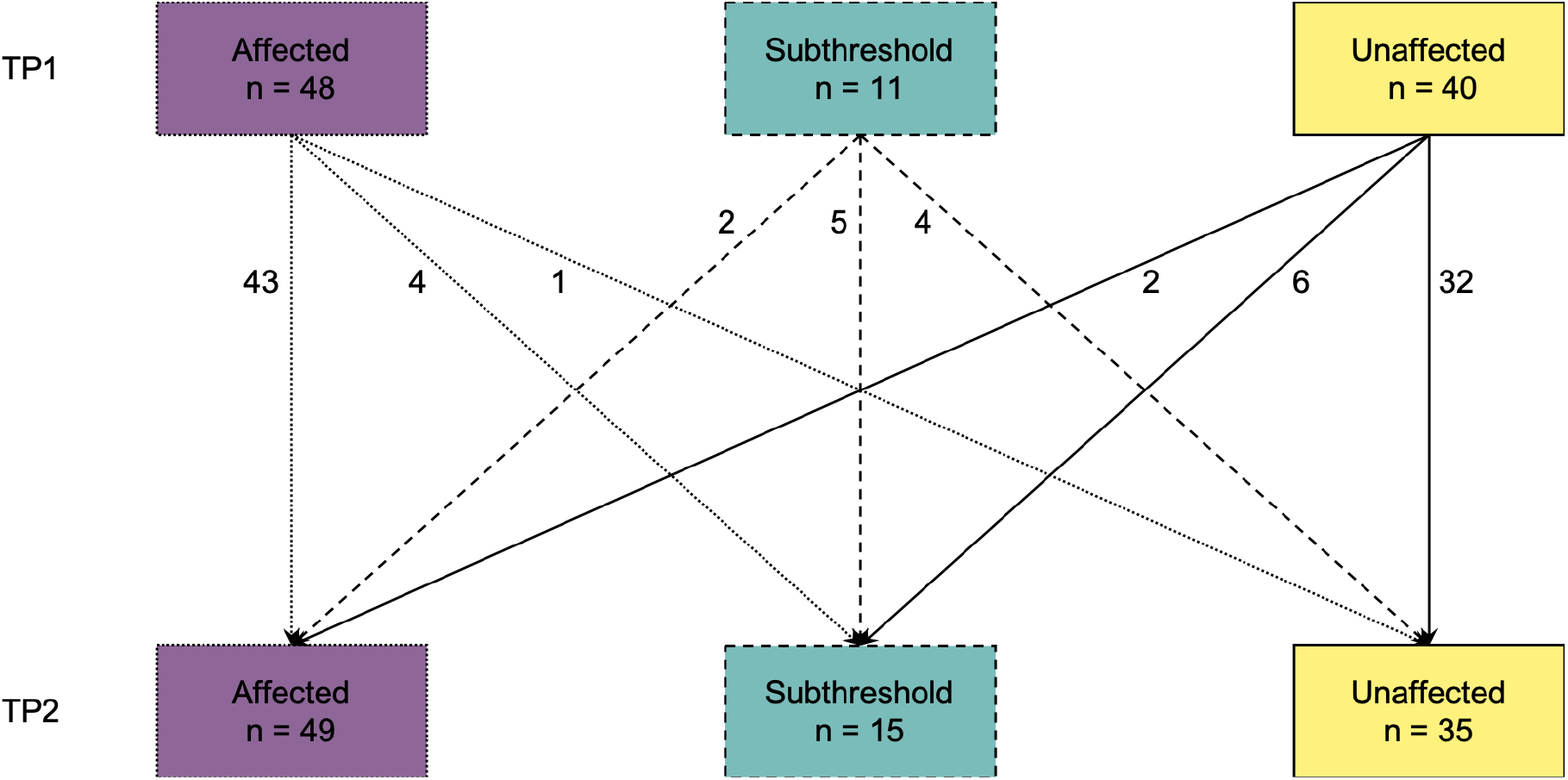
Changes in the diagnostic make-up of our study sample (N=99). Although the total number of participants remained constant, the number of individuals in each diagnosis group changed between time-point 1 (TP1) and time-point 2 (TP2).

**Figure S4.**
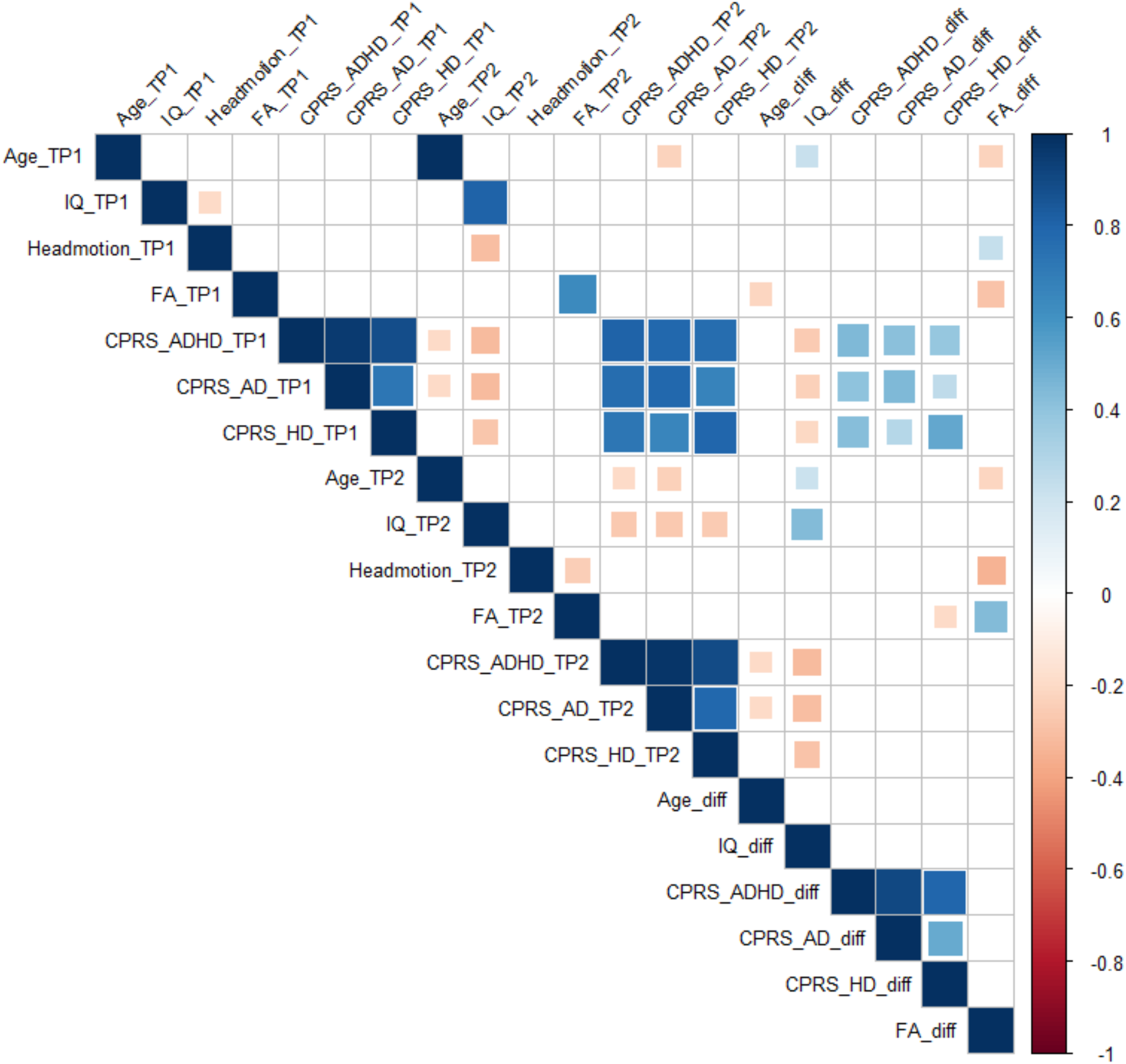
Spearman correlation matrix of independent and dependent variables, as well as covariates. These correlation tests were performed before the main analyses. The color intensity of each box indicates the magnitude of the correlation. Positive correlations are presented in blue and negative correlations in red. The size of the box indicates its significance, with significant correlations filling each square completely.

**Figure S5.**
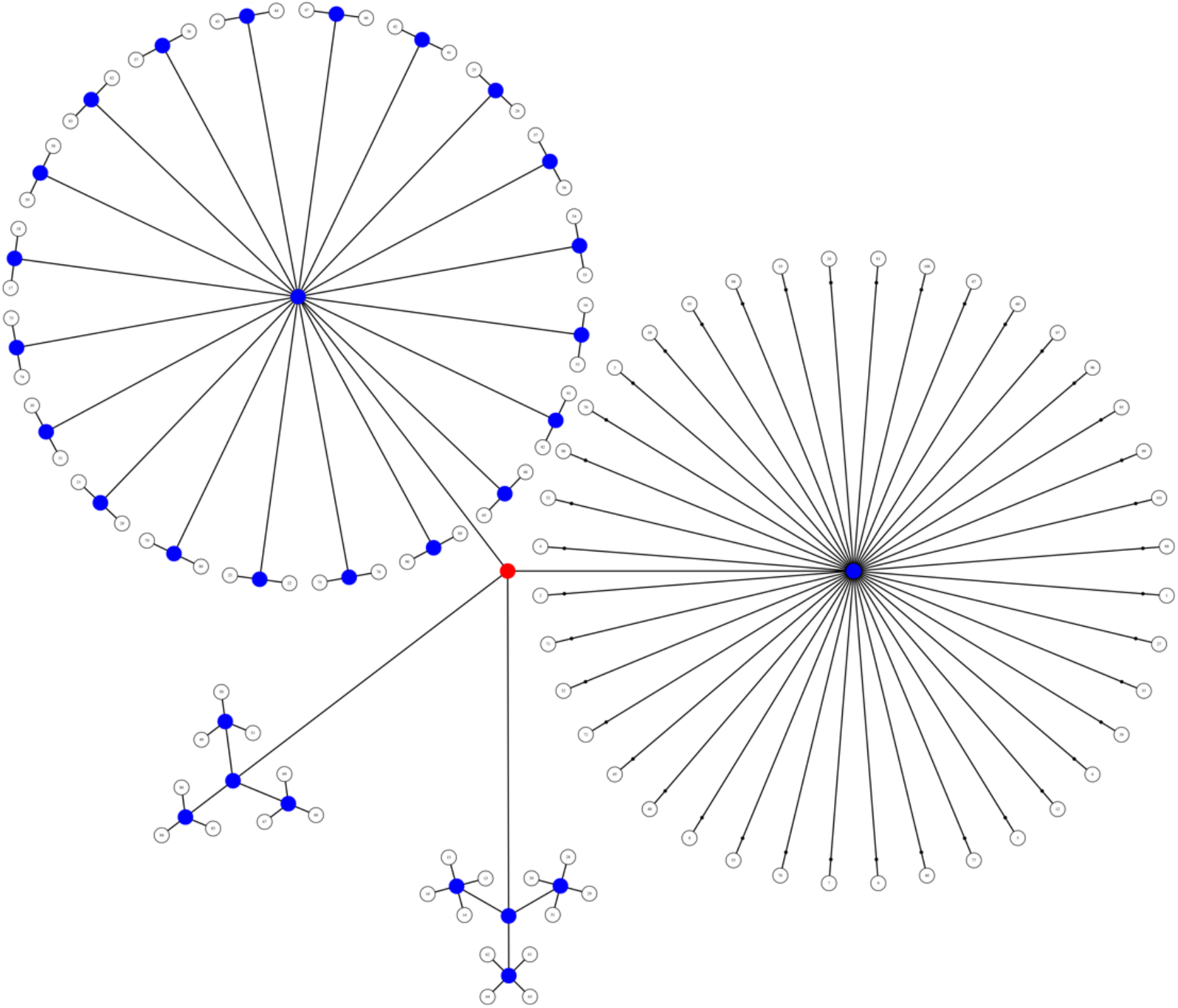
Visual representation of the multi-level notation of family structure in our sample. The four groups represent the size of each family: 3 families of 3 children, 3 families of 4 children, 20 families of 2 children, and 40 families of 1 child in the study. We depict the levels as branches from the central red node, akin to a tree in which the most peripheral elements (leaves) represent the observations. The nodes from which the branches depart either allow (blue) or do not allow (red) permutations.

**Figure S6.**
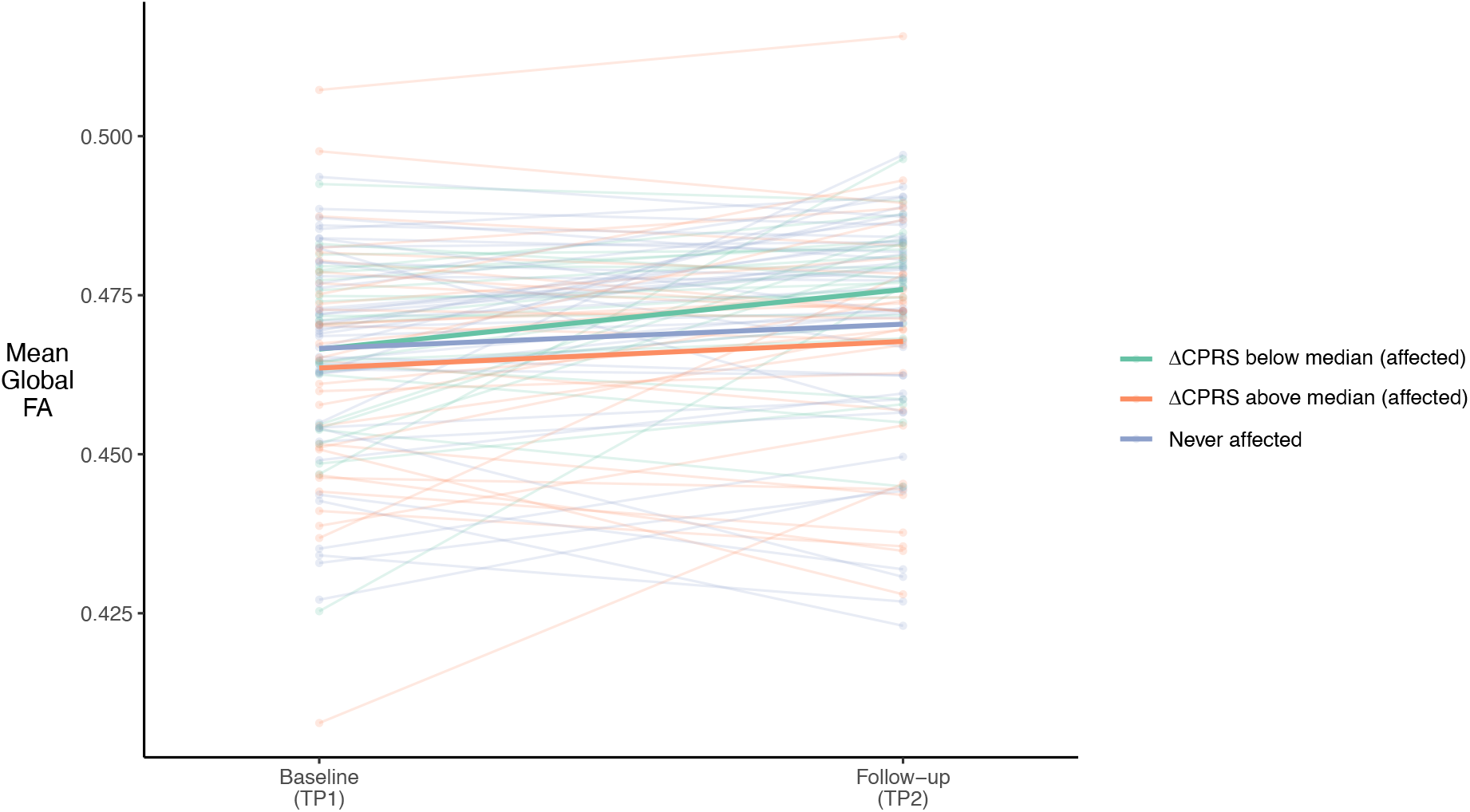
Line plot illustrating individual changes in average whole-brain fractional anisotropy (FA) from baseline to follow-up, grouped by never affected participants (blue) versus affected participants (including subthreshold). Here, affected individuals are further split along the median into two groups according to whether their change in combined Conners Parent Rating Scale score (ΔCPRS) was above (orange) or below (green) the affected sample’s median of that change.

**Figure S7.**
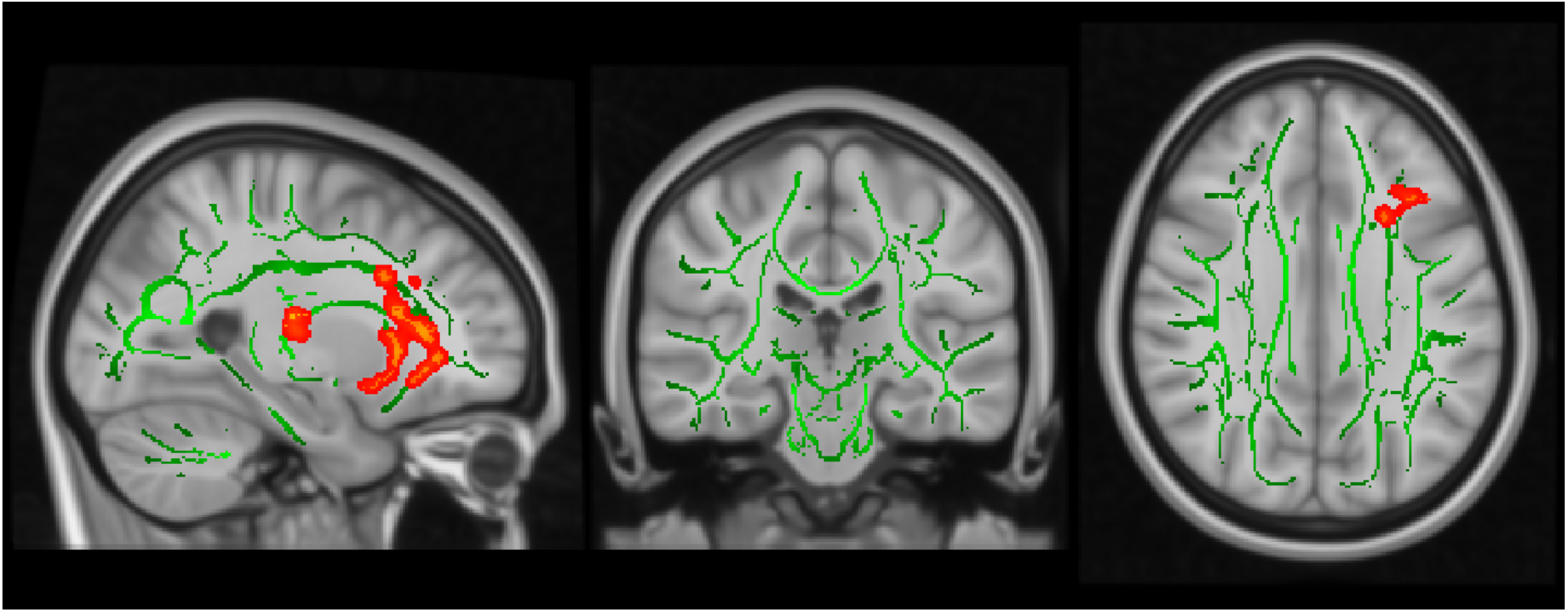
Exploratory dimensional TBSS analyses showing significant associations (red-yellow) between FA values and the CPRS scores over time. The mean FA skeleton across all subjects (green) was overlain on the MNI template image for presentation (x=−25, y=−25, z=31). Results were thickened for visualization (FSL “tbss_fill”) and presented here in radiological convention from sagittal, coronal, and axial perspectives, respectively. A more negative change in FA (i.e. more isotropic diffusion) over time was associated with more HI symptom remission in ten clusters spread over six WM tracts. See Table S2 for cluster statistics and locations.

## Notes

### Competing Interest Statement

The authors have declared no competing interest.

